# Comparison of muscle activity, strength and balance, before and after a 6-month training using the FIFA11+ program (part 2)

**DOI:** 10.1101/540492

**Authors:** Takeshi Oshima, Junsuke Nakase, Anri Inaki, Takafumi Mochizuki, Yasushi Takata, Kengo Shimozaki, Seigo Kinuya, Hiroyuki Tsuchiya

**Author notes:** Corresponding author: Junsuke Nakase, M.D., Ph.D., Department of Orthopaedic Surgery, Graduate School of Medical Science, Kanazawa University 13-1 Takara-machi, Kanazawa 920-8641, Japan.

## Abstract

**Purpose:** Sports injuries can significantly impact an athlete’s career, as well as impose a high financial burden on teams. Therefore, the prevention of sports injuries is an essential aspect of sports medicine. To evaluate the effects of a 6-month training period, using part 2 of the FIFA11+, on the activation and strength of core and lower limb muscles, and on static and dynamic balance performance.

**Materials and Methods:** Eight college male soccer players, 20.4±0.5 years old, completed the FIFA11+ program (part 2) at least 3x per week for 6 months. The following variables were measured, before and after the 6-month training program: activity of more than 30 muscles (with a focus on core and lower limb muscles), measured using the standardized uptake values of 18F-fluorodeoxyglucose (FDG) on positron emission tomography (PET-CT); isokinetic strength of the knee flexor and extensor and hip abductor muscles, measured at 60°/s; static balance over a 60-s period, measured using a Gravicorder; and dynamic balance, measured using the Star Excursion Balance Test.

**Results:** Training improved activity levels of core (obliquus externus abdominis and erector spinae) and lower limb (tibialis anterior of the both legs) muscles (p≤0.03), corrected the between-limb difference in activation of the semimembranosus and improved dynamic balance, with a greater training effect on the non-dominant limb (p≤0.02). Training also improved knee flexor force of the non-dominant lower limb (p=0.02).

**Conclusion:** Routine performance of the FIFA11+ (part 2) program can improve activation of core and lower limb muscles, with a concomitant improvement in dynamic balance.

## Introduction

Sports injuries can significantly impact an athlete’s career and the financial aspect [1]. Therefore, the prevention of sports injuries has received increasing attention in sports medicine. Generally, sports injury prevention programs include some combination of plyometric, balance and agility exercises, and have been reported to be effective in decreasing the incidence of injuries, regardless of sport activity level, sex and age [2,3]. ‘FIFA11+’ is one of the most effective prevention programs, which the Fédération Internationale de Football Association (FIFA) Medical and Assessment Research Center (F-MARC) has developed. The FIFA11+ consists of three parts: basic running (part 1); 3 levels of difficulty of 6 exercises aiming to increase strength (core and lower limbs), balance, muscle control (plyometrics), and core stability (part 2); and running such as straight line running, or cutting activities (part 3).

Improvement and evaluation of the effectiveness of an injury prevention program requires assessment of not only the change in the incidence rate of injury, but also the short- and long-term effects of the training on modifying muscle activity patterns and improving, strength and balance.

To evaluate the muscle activities, electromyography (EMG) has principally been used. However, EMG can only provide information on the activation of superficial muscles, and not of deep muscles of the trunk and limbs, is limited with regard to the number of muscles that can be assessed simultaneously (namely those on which superficial sensors can be placed) and requires equipment to be attached to the body, which is difficult during performance of sports activities.

Previous studies have used whole-body positron emission tomography - computed tomography (PET-CT) to quantify the change in muscle activity after performing the FIFA11+ (part 2) program, with glucose uptake in skeletal muscles being used as a proxy measure of the level of muscle activity [4,5]. Unlike EMG, PET-CT provides a non-invasive observation of the activity of muscles throughout the body, simultaneously, with the possibility of 3-dimensional (3D) image reconstruction. As active muscle cells exhibit increased glucose uptake, the use of ^18^F-fluorodeoxyglucose (FDG), a deoxy analog of glucose, permits the observation of glucose metabolism of the skeletal muscles throughout the body. However, unlike glucose, FDG does not continue along the usual glycolytic pathway but, rather, accumulates within exercising muscle tissue. This metabolic trapping process forms the basis of FDG-PET. The accumulation of FDG in muscle provides a parameter of glucose intake by muscles and, therefore, of the intensity of muscle activity [6]. To our knowledge, FDG-PET is the only method that can provide a reliable cumulative index of muscle activity for between-muscle comparison, which is invaluable for the assessment of sports injuries. As such, FDG-PET would be effective to measure the effects of the FIFA11+ program.

In recent years, various benefits of the entire FIFA11+ program have been reported, including a reduction in the incidence of sports injuries and improvement in the neuromuscular control and strength of flexor muscles [7,8]. A review of the studies reporting on the acute or chronic effects of the FIFA11+ on performance and physiological measures among football players, an intervention period of 9-10 weeks yielded positive effects [9]. However, the effect of a long-term, routine, performance of the FIFA11+ program on the metabolism of skeletal muscles remains to be clearly defined. Therefore, the aim of our study was to investigate the change in muscle activity, and muscle strength and dynamic balance, after performing part 2 of the FIFA11+ for 6 months. The a priori hypothesis was that the FIFA11+ (part 2) program would be effective in increasing the activity of core and lower limb muscles, improve muscle strength of the lower limb and improve static and dynamic balance.

## Materials and Methods

Our study group was formed of 8 collegiate male soccer players. All participants were considered to be healthy, based on their medical history and physical examination, and none were taking medication at the time of the study. All participants provided informed consent, and the study was approved by our Institutional Ethics Review Board.

Participants were asked to avoid strenuous physical activity for at least one day prior to testing, and to refrain from eating and drinking for at least 6 h before testing. PET-CT images were obtained as per previously described methods [4,5]. After obtaining baseline (pre-training) PET-CT images, participants completed the training protocol, consisting of completing part 2 of the FIFA11+ program, ≥3 times per week, for 6 consecutive months. PET-CT images were obtained at the end of the training period, using the same protocol as at baseline.

Regions of interest (ROI) on the images were manually segmented in 30 skeletal muscles, located in 5 areas of the body: trunk, pelvis, thigh, lower leg, and the foot. (Table 1) All ROIs were identified by one experienced nuclear medicine specialist, who was blinded from all other results, from the plain CT images obtained concurrently. The standardized uptake value (SUV) of FDG was calculated by overlapping the defined ROI and fusion PET-CT images to outline the area of muscles, being careful to not include large vessels. The FDG uptake was normalized to the unit volume of muscle as follows: {mean ROI count (counts per second/pixel) × calibration factor (counts per second /Bq)}/{injected dose (Bq)/body weight (g)}. ROIs were defined for the skeletal muscles previously described, bilaterally, and the mean SUV was compared for the dominant and non-dominant side of the body (where the dominant side was identified by asking participants which leg they used to kick a ball). The mean SUV for the trunk was calculated as follows: ([left mean SUV × left muscle area] + [right mean SUV × right muscle area])/(left muscle area + right muscle area). FDG accumulation was compared between the pre- and post-training PET-CT examinations.

**Table 1:** Mean SUVs during pre- and post-training. SUV, standardized uptake value

| Body area | Muscles | Pre-training SUV |  | Post-training SUV |  |
| --- | --- | --- | --- | --- | --- |
|  |  | Dominant leg | Non-dominant leg | Dominant leg | Non-dominant leg |
| Trunk | Rectus abdominis | 0.90±0.35 |  | 1.03±0.40 |  |
|  | Obliquus externus abdominis | 0.75±0.26 |  | 1.06±0.38 |  |
|  | Obliquus internus abdominis | 0.69±0.14 |  | 0.76±0.81 |  |
|  | Transversus abdominis | 0.62±0.08 |  | 0.59±0.14 |  |
|  | Psoas major | 0.94±0.31 |  | 0.87±0.20 |  |
|  | Quadratus lumborum | 0.85±0.33 |  | 0.99±0.32 |  |
|  | Erector spinae | 0.67±0.16 |  | 0.80±0.31 |  |
| Pelvis | Gluteus maximus | 1.54±0.78 | 1.25±0.78 | 1.27±0.29 | 1.17±0.29 |
|  | Gluteus medius | 2.18±1.17 | 2.29±0.82 | 2.79±1.93 | 2.46±0.31 |
|  | Gluteus minimus | 3.13±0.60 | 3.51±0.83 | 3.51±1.20 | 3.34±0.98 |
|  | Piriformis | 2.95±1.70 | 2.37±0.88 | 2.13±0.74 | 1.99±0.65 |
| Thigh | Quadriceps femoris | 1.00±0.31 | 1.01±0.38 | 1.00±0.30 | 0.92±0.19 |
|  | Sartorius | 0.76±0.21 | 0.77±0.20 | 0.91±0.32 | 0.89±0.27 |
|  | Gracilis | 1.16±0.43 | 1.14±0.40 | 1.26±0.43 | 1.32±0.64 |
|  | Semimembranosus | 0.74±0.14 | 0.59±0.10 | 0.62±0.08 | 0.60±0.12 |
|  | Semitendinosus | 1.19±0.31 | 1.09±0.30 | 1.41±0.48 | 1.23±0.52 |
|  | Biceps femoris | 0.59±0.08 | 0.63±0.10 | 0.61±0.10 | 0.62±0.13 |
|  | Adductor complex | 0.81±0.18 | 0.82±0.18 | 0.83±0.14 | 0.76±0.15 |
| Lower leg | Tibialis anterior | 1.06±0.59 | 1.00±0.36 | 1.53±0.86 | 1.44±0.66 |
|  | Flexor digitorum longus | 1.28±0.26 | 1.20±0.35 | 0.90±0.35 | 0.96±0.35 |
|  | Tibialis posterior | 1.34±0.84 | 1.37±0.79 | 1.05±0.30 | 1.13±0.51 |
|  | Flexor hallucis longus | 1.23±0.13 | 1.30±0.58 | 1.10±0.29 | 1.18±0.49 |
|  | Peroneus | 1.50±0.89 | 1.32±0.36 | 1.13±0.61 | 1.21±0.52 |
|  | Triceps surae | 1.39±0.40 | 1.24±0.25 | 0.88±0.19 | 0.86±0.18 |
| Foot | Abductor hallucis | 1.36±0.42 | 1.45±0.44 | 1.31±0.61 | 1.17±0.37 |
|  | Quadratus plantae | 1.06±0.20 | 1.29±0.43 | 1.05±0.38 | 1.00±0.34 |
|  | Flexor digitorum brevis | 1.43±0.42 | 1.61±0.58 | 1.68±1.49 | 1.49±0.67 |
|  | Abductor digiti minimi | 1.13±0.44 | 1.45±0.55 | 1.15±0.50 | 1.31±0.78 |
|  | Flexor hallucis brevis | 2.12±0.65 | 2.24±0.57 | 2.27±0.83 | 2.12±0.90 |
|  | Interosseous | 2.02±0.61 | 2.20±1.11 | 2.38±1.50 | 2.20±1.31 |

Balance and muscle strength testing was performed by experienced physical therapists. These assessments were performed one week after the PET analysis, with balance tests preceding strength tests to avoid effects of muscle fatigue.

Static balance was measured using a Gravicorder. Postural sway, for 60 s, at a sampling rate of 20-Hz sampling, under the following conditions: two-leg stance with eyes open and then with eyes closed; single leg (dominant) standing with eyes open; and single leg (non-dominant) standing with eyes open. All measurements were obtained in bare feet, using the center of the force platform as a reference point. Two variables of balance were measured, the locus length per time (LG), providing a measure of attitude control, and the environmental area (AR), providing a measure of equilibrium control [10]. These two parameters have been used to assess dizziness and equilibrium disorders and, more recently, to quantify balance effects on anterior cruciate ligament injury [11]. All balance measurements were repeated twice, with a 1-min rest between measurements; data from the second measurement, which has been reported to be more accurate [12], used for analysis.

Dynamic postural control was evaluated using the Star Excursion Balance Test (SEBT). Participants were asked to reach as far as possible along the designated line for each of the following 8 directions: anterolateral, anterior, anteromedial, medial, posteromedial, posterior, posterolateral, and lateral. (Fig 1) The test was performed twice, once in a clockwise direction (reaching with the right leg) and once in a counterclockwise direction (reaching with the left leg). The average of the length of three reaches performed in each direction was used for analysis, with the distance normalized to the length of the leg (measured from the anterior superior iliac spine to the distal tip of the medial malleolus). The greater the normalized length of excursion, the better the dynamic balance.

**Fig 1:**
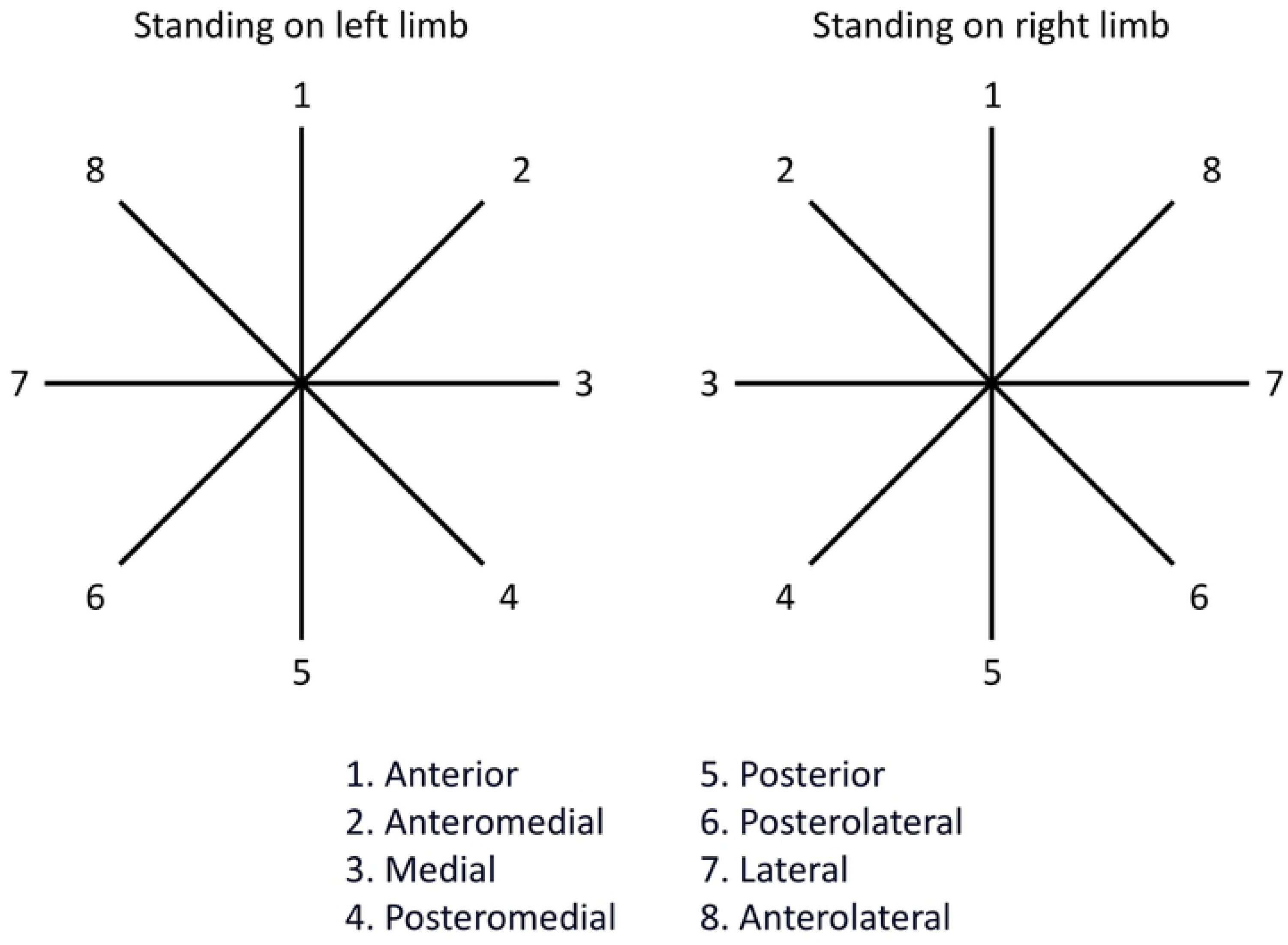
Reaching directions on the Star Excursion Balance Test.

The maximal knee flexion and extension muscle strength and the maximal isokinetic hip abductor strength were tested using an isokinetic Biodex system. After 10-min warm-up, knee strength measurements were obtained in the sitting position, and the hip abductor strength was obtained in a sidelying position. For the strength testing, participants were asked to move the knee and hip at full force, with 3 trials performed for each direction at a speed of 60°/s. For all strength measurements, the average value of the 3 trials was used for analysis, and values were normalized to body weight (pre- and post-training) for between-subject comparisons. The ratio of the strong-to-weaker leg was calculated as an index of between-limb strength imbalance, converted to a percentage difference, using a previously described method based on log-transformed raw data, followed by back transformation [13].

All statistical analyses were performed using Stata for Mac Version 15 (Stata Statistical Software 2017; Stata Corp LLC, College Station, TX, USA). All data are presented as mean (SD). The Shapiro-Wilk test was used to evaluate the normality of distribution. Wilcoxon signed rank test was used to evaluate differences in the mean SUV and static balance, before and after training, with a paired t-test used to evaluate the differences in muscle strength and dynamic balance. The minimum significance level was set at P < 0.05. The sample size was confirmed using a power analysis of 0.8, with an α value of 0.05 and effect size of 1.0.

## Results

The relevant characteristics of the participants at pre-training are as follows: age; 20.4±0.5 years old, height; 175.4±6.2 cm, weight; 68.6±5.1 kg, 22.3±1.3 kg/m^2^, and the leg length; 89.4±3.8 cm. After training, the weight was 70.1±4.6 kg (p= 0.246) and the body mass index was 22.8±0.8 kg/m^2^ (p= 0.250), and there was no significant difference pre- and post-training.

Representative whole-body PET images, pre- and post-training, are shown in Fig 2 with the mean SUVs reported in Table 1. A significant pre- to post-training increase in the mean SUV was identified for two core muscles, the obliquus externus abdominis (0.75±0.26 *versus* 1.06±0.38, respectively, p= 0.036) and erector spinae (0.67±0.16 *versus* 0.80±0.31, respectively, p=0.025). The pre- to post-training significant change in the mean SUVs for the muscles of the dominant and non-dominant lower limbs, from the pelvis to the foot, are detected. For the dominant lower limb, the mean SUV increased for the tibialis anterior (1.06±0.59 *versus* 1.53 ±0.86, respectively, p=0.017) and decreased for the triceps surae (1.39±0.40 *versus* 0.88±0.19, respectively, p=0.017). A similar result was identified for the non-dominant lower limb, with an increase in the tibialis anterior (1.00±0.36 *versus* 1.44±0.66, respectively, p=0.025) and a decrease for the triceps surae (1.24±0.25 *versus* 0.86±0.18, respectively, p=0.025). The significant side-to-side difference of SUV was detected in semimembranosus in pre-training. Pre-training, the mean SUV of the semimembranosus muscle was higher for the dominant than non-dominant lower limb (0.74±0.14 *versus* 0.59±0.10, respectively, p=0.012). Of note, no significant difference in the activation of the semimembranosus between the dominant and non-dominant side was observed after training.

**Fig 2:**
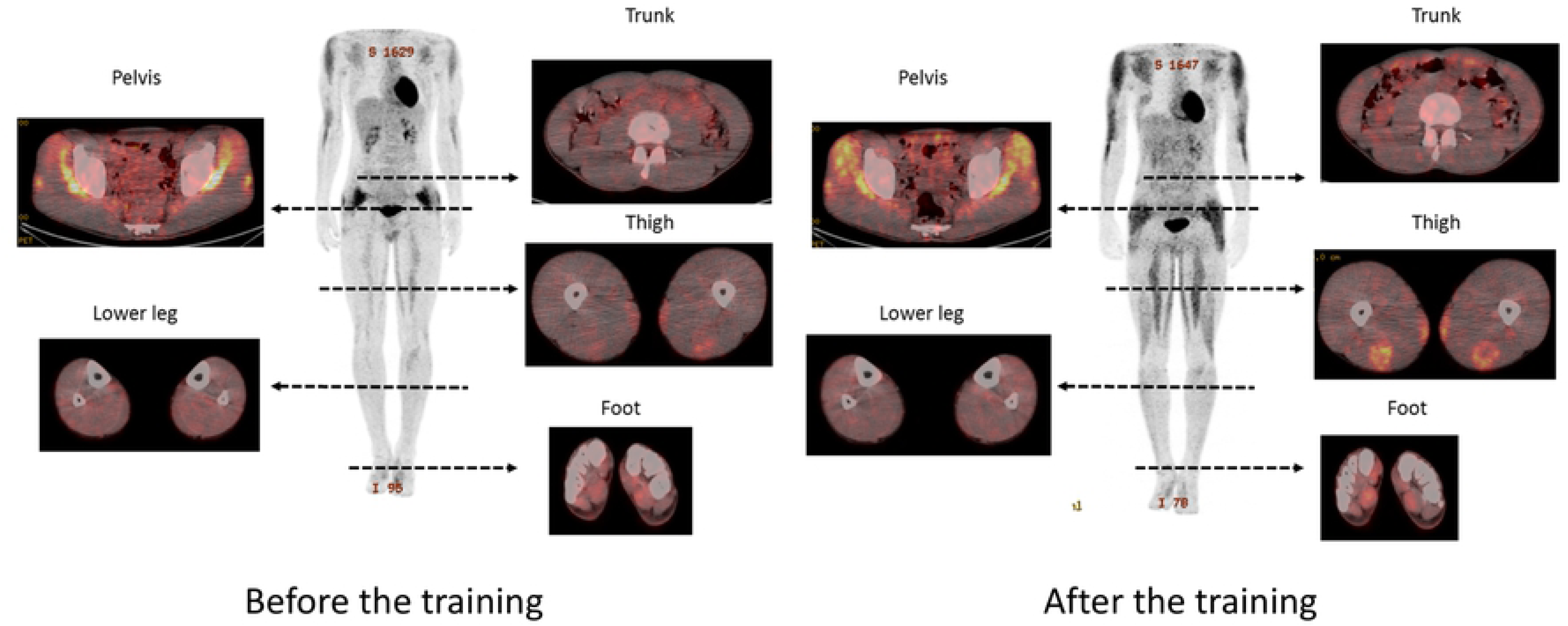
Representative whole-body positron emission tomography images acquired before (left) and after (right) routine performance of the FIFA11+.

The mean LG and AR values are reported in Table 2, with no significant difference between pre- and post-training values. The reach distance along the 8 directions for the dominant and non-dominant lower limbs are reported in the Table 3. For the dominant leg, the standing reach distance increased significantly, pre- to post-training, in the anterior-lateral direction (70.9±9.2 cm to 74.9±9.6 cm, p=0.023). A greater improvement in dynamic balance on the non-dominant leg was observed, with an increase in the reach distance across multiple directions, as follows: medial (103.6±6.0 cm to 107.9±7.6 cm, p=0.002); posterior-medial (111.4±6.0 cm to 115.3±6.6 cm, p=0.030); and posterior (114.0±4.5 cm to 119.1±7.9 cm, p=0.022). The pre- to post-training changes in muscle strength are reported in Table 4. For the non-dominant leg, knee flexion force increased from 1.24±0.15 Nm/kg to 1.39±0.14 Nm/kg (p=0.023). No effects of training on knee extensor and hip abductor strength were noted, nor on the hamstring-to-quadriceps ratio or between-limb imbalance index (Table 5).

**Table 2:** Static balance parameter, pre- and post-training.

|  | conditions | pre-training | post-training | Confidence interval | Effect size | P value |
| --- | --- | --- | --- | --- | --- | --- |
| LG (cm/s) | Two-leg, eyes opened | 0.94±0.11 | 1.05±0.14 | [-0.04, 0.27] | 0.904 | 0.12 |
|  | Two-leg, eyes closed | 1.41±0.28 | 1.27±0.23 | [-0.39, 0.11] | 0.533 | 0.36 |
|  | Dominant leg, eyes opened | 3.93±0.61 | 3.96±0.67 | [-0.44, 0.51] | 0.052 | 0.78 |
|  | Non-dominant leg, eyes opened | 3.88±0.68 | 4.03±0.31 | [-0.48, 0.79] | 0.195 | 0.40 |
| AR (cm<sup>2</sup>) | Two-leg, eyes opened | 1.69±0.64 | 1.81±0.40 | [-0.44, 0.69] | 0.181 | 0.48 |
|  | Two-leg, eyes closed | 2.65±0.37 | 2.46±0.82 | [-1.01, 0.62] | 0.198 | 0.78 |
|  | Dominant leg, eyes opened | 6.84±2.77 | 7.24±1.56 | [-2.30, 3.11] | 0.124 | 0.78 |
|  | Non-dominant leg, eyes opened | 6.91±2.66 | 7.45±1.99 | [-2.10, 3.18] | 0.177 | 0.67 |

**Table 3:**
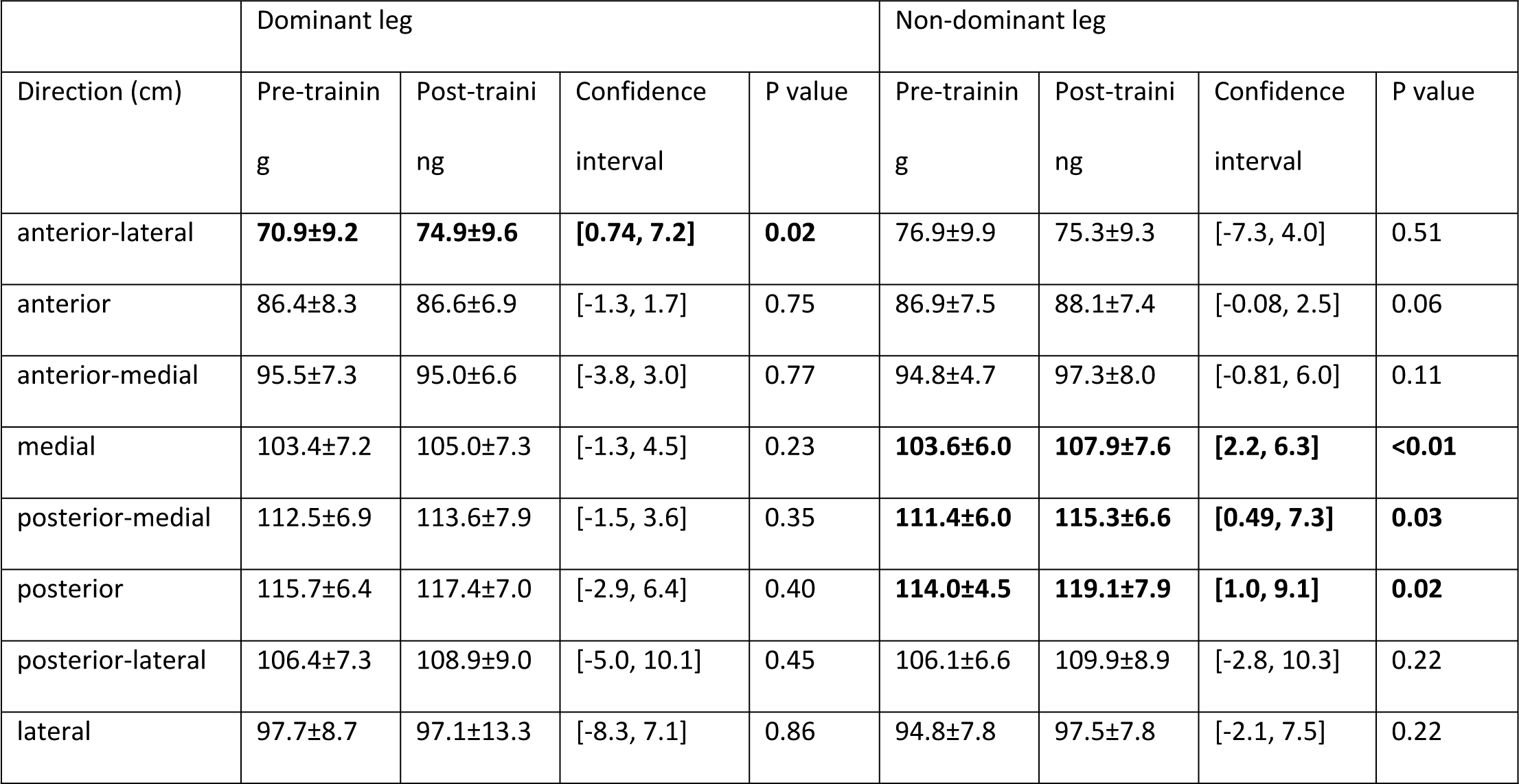
Reach distance (excursion distance/leg length×100) by balance condition and direction of reach, pre- and post-training.

**Table 4:** Pre- to post-training change in lower limb muscle strength.

|  | pre-training | post-training | Confidence interval | Effect size | P value |
| --- | --- | --- | --- | --- | --- |
| Knee flexor (%) | 17.0±22.3 | 6.5±4.9 | [-28.5, 7.52] | 0.487 | 0.22 |
| Knee extensor (%) | 12.5±8.2 | 12.3±8.6 | [-10.7, 10.3] | 0.016 | 0.96 |
| Hip abductor (%) | 9.9±7.9 | 6.7±5.0 | [-10.5, 4.0] | 0.373 | 0.33 |
| H/Q ratio (%) | 17.5±5.8 | 10.6±8.4 | [-16.4, 2.7] | 0.596 | 0.14 |

**Table 5:** Between-limb muscle strength imbalance, pre- and post-training.

|  | Dominant leg |  |  |  |  | Non-dominant leg |  |  |  |  |
| --- | --- | --- | --- | --- | --- | --- | --- | --- | --- | --- |
|  | pre-training | post-training | Confidence interval | Effect size | P value | pre-training | post-training | Confidence interval | Effect size | P value |
| Knee flexor (Nm/kg) | 2.90±0.09 | 2.99±0.18 | [-0.47, 6.44] | 0.134 | 0.72 | 2.78±0.17 | 2.98±0.13 | [-0.10, 0.50] | 0.550 | 0.16 |
| Knee extensor (Nm/kg) | 1.31±0.20 | 1.41±0.21 | [-0.15, 0.34] | 0.338 | 0.37 | **1.24±0.15** | **1.39±0.14** | **[0.03, 0.28]** | **1.026** | **0.02** |
| Hip abductor (Nm/kg) | 2.41±0.41 | 2.40±0.35 | [-0.41, 0.38] | 0.031 | 0.21 | 2.35±0.37 | 2.52±0.43 | [-0.05, 0.39] | 0.634 | 0.55 |
| H/Q ratio (%) | 0.45±0.05 | 0.48±0.07 | [-0.02, 0.08] | 0.454 | 0.16 | 0.46±0.09 | 0.48±0.09 | [-0.05, 0.09] | 0.220 | 0.78 |

## Discussion

Our results indicate an increase in the activation of various skeletal muscles of the core and lower limbs after a 6-month training using part 2 of the FIFA 11+ program, measured as an increase in the uptake of glucose: obliquus externus abdominis, erector spinae and tibialis anterior. Of note was the decrease in the glucose uptake of the triceps surae, as glucose uptake increased in the tibialis anterior. We also noted an improvement in the imbalance of the glucose uptake in the semimembranosus, between the dominant and non-dominant, after training. From a functional perspective, training produced a greater improvement in dynamic balance on the non-dominant than dominant lower limb. To our knowledge, this is the first study to report changes in muscle activities, associated with improvements in balance and muscle strength, after long-term training using the part 2 of the FIFA 11+ program. Observed changes in glucose uptake, balance and strength would, therefore, be the key mechanisms explaining the association between the FIFA 11+ and a decrease in sports-related injuries.

Glucose enters the muscle cell by facilitated diffusion via the glucose transporter-4 (GLUT4), with exercise stimulating an increase in the expression of GLUT4 in skeletal muscles, as shown by the findings of Reichkendler et al. after an 11-week program of daily moderate- and high-dose aerobic exercise [14]. Similarly, an increase in GLUT4 levels in skeletal muscles is a key adaptation to regular exercise training [15]. Thus, FDG accumulation in the muscle can be used as a measure of the change in glucose uptake with training, as well as providing a proxy measure of muscle activity [16].

By comparing the change in FDG accumulation of each muscle from pre- to post-training, we demonstrated that routine training using part 2 of the FIFA11+ program improved muscle metabolism and activation as well as the previous studies [4,5]. These adaptations are important when we consider the positive effects of core and lower limb strength on balance. Kaji et al. reported on the improvement in two-leg standing balance with eyes closed after performing the FIFA11+ program (p=0.005) [17]. Granacher et al. reported on the improvement in the Functional Reach test (p<0.05) after performing a core stability training program which increased the strength of the trunk flexors (p<0.001), extensors (p<0.001) and lateral flexors (p<0.001) [18]. In the same way, Imai et al. reported immediate improvements in the posteromedial (p<0.001) and posterolateral directions (p=0.002) of the SEBT after the trunk stabilization exercises [19]. Considering the effect of core stability on balance, Willson et al. suggested that appropriate core strength training could reduce sports-related injuries [20]. We reported similar findings, showing an increase in the mean SUVs of the obliquus externus abdominis and erector spinae muscles after training, with a concomitant improvement in dynamic balance, indicative of the effectiveness of the part 2 FIFA11+ program in improving core strength.

With regard to lower limb muscle activation, Day et al. reported an increase activity of the tibialis anterior during active swaying (compared to static standing), which they associated to the higher proprioceptive demands of balancing under more challenging sensory conditions and the proprioceptive role of the tibialis anterior [21]. Similarly, Earl et al. reported an increase in the general activity of lower limb muscles (vastus medialis obliquus, vastus lateralis, medial hamstring, biceps femoris, and tibialis anterior) during the SEBT (p<0.05), with the exception of the triceps surae muscles (p=0.08) [22]. We demonstrated comparable findings post-training, supporting the effectiveness of the FIFA11+ (part 2) in improving balance [8].

The balance of muscle strength is also an important component with regard to injury prevention. For the lower limb, the hamstring-to-quadriceps ratio is an important risk factor of injury [23]. Previous studies have reported on the effectiveness of the complete FIFA 11+ program in improving knee flexor strength and, thus, the hamstring-to-quadriceps ratio [7]. Between-limb strength imbalances might also be an important risk factor for lower leg injury [24]. A prospective study provided evidence that a between-limb imbalance in eccentric knee flexor strength increase in the risk of hamstring injuries [25]. Although we reported the effectiveness of the training in eliminating the higher SUV of the semimembranosus muscles in the dominant than non-dominant lower limb observed pre-training, we did not identify a significant between-limb imbalance among our study participants, either pre- or post-training. Therefore, we cannot stipulate if the FIFA11+ (part 2) program is effective to improve lower limb muscle imbalances, although the results of our SUV analysis indicate that the program would likely be of benefit in this regard.

Overall, our findings are consistent with previous reports on the effectiveness of performing the complete FIFA 11+ program in improving balance and muscle strength, which lowered the incidence of sports injuries [7-9]. The methods we used, and PET-CT in particular, could be useful for evaluating the effectiveness of training programs and identifying the underlying pathways.

We note the following limitations of our study. First, the FDG-PET method accounts only for muscle glucose uptake. Other substrates, such as free fatty acids, muscle glycogen and lactate, are also metabolized in active muscle cells. That being said, studies have confirmed that glucose oxidation increases with exercise intensity and increases in glucose uptake, to some extent, increases in proportion with glycogen utilization as exercise intensity increases. A second limitation was the method we used to define the ROI. Since FDG uptake was measured at an arbitrary site on the target muscle, it did not reflect the uptake of glucose for the entire muscle. Lastly, taking into consideration the ethical dilemma of radiation exposure with CT (even though the amount of FDG was <10% of normal PET examination), our study sample was small and we did not include a control group. Our use of PET-CT was consistent with the aim of our study to confirm that improvements in muscle metabolism was an important underlying pathway for the previously reported effectiveness of the FIFA11+ training program. In this sense, our use of PET-CT and our findings of an increase in glucose uptake post-training are novel.

## Conclusions

Routinely performing part 2 of the FIFA11+ program for 6 months increased glucose uptake, related to muscle activity, of the obliquus externus abdominis, erector spinae and tibialis anterior, while decreasing activation of the triceps surae. The training program also improved knee flexor strength and dynamic balance, with no effect identified on static balance. We speculate that these improvements could be beneficial in lower the risk of sports-related injuries.

## Acknowledgments

The authors would like to express their appreciation for the outstanding efforts positive attitude of the participants. In addition, they are extremely grateful for the technical assistance with measurement provided by physical therapists of our hospital. No financial assistance was received for the project.

## Author Contributions

Conceptualization: Takeshi Oshima, Junsuke Nakase.

Data curation: Takeshi Oshima, Anri Inaki.

Formal analysis: Takeshi Oshima, Anri Inaki, Takafumi Mochizuki.

Investigations: Takeshi Oshima, Yasushi Takata, Kengo Shimozaki.

Methodology: Takeshi Oshima, Anri Inaki, Takafumi Mochizuki, Junsuke Nakase.

Project administration: Seigo Kinuya, Hiroyuki Tsuchiya.

Resources: Takeshi Oshima, Yasushi Takata, Kengo Shimozaki, Junsuke Nakase.

Supervision: Seigo Kinuya, Hiroyuki Tsuchiya.

Validation: Takeshi Oshima, Anri Inaki, Takafumi Mochizuki.

Writing-original paper: Takeshi Oshima.

Writing-review and editing: Junsuke Nakase, Seigo Kinuya, Hiroyuki Tsuchiya.

## Conflict of interest statement

This research didn’t receive grants from any funding agency in the public, commercial or not-for-profit sectors.

## Ethical approval

This study was approved by the ethics committee of Kanazawa University (approval number: 1286).

